# Comparative genomics of *Staphylococcus capitis* reveals determinants of speciation

**DOI:** 10.1101/2022.02.11.480064

**Authors:** Charlotte E. Chong, Rebecca J. Bengtsson, Malcolm J. Horsburgh

## Abstract

*Staphylococcus capitis* is primarily described as a human skin commensal but is now emergent as an opportunistic pathogen isolated from bloodstream and prosthetic joint infections, and neonatal intensive care unit (NICU) associated sepsis. We used comparative genomic analyses of *S. capitis* to provide new insights of commensal scalp isolates from varying skin states, and to expand our current knowledge of the species populations (scalp isolates, *n* = 59, contextual isolates, *n* = 127). A highly recombinogenic population structure was revealed, with genomes including the presence of a range of previously described staphylococcal virulence factors, cell wall-associated proteins, and two-component systems. Genomic differences between the two described *S. capitis* subspecies were explored and reveal determinants associated exclusively with each. The subspecies *ureolyticus* was distinguished from subspecies *capitis* by differences in antimicrobial resistance genes and gene clusters linked to survival on the skin. This study will aid further research into classification of *S. capitis* and virulence linked phylogroups that is important to monitor the spread and evolution of *S. capitis.*

## Introduction

*Staphylococcus capitis* was first isolated from healthy human skin in 1975 and classified as a coagulase-negative *Staphylococcus* species (CoNS) (1). *S. capitis* is frequently found on the human scalp and the forehead, as it thrives in lipid-rich areas where sebaceous glands are abundant and was more recently associated with dandruff presenting scalps (1–4). *S. capitis* can be divided into two subspecies, *capitis* and *ureolyticus,* based on differences in their urease production and maltose fermentation (2). Current literature has additionally sought to characterise the two *S. capitis* subspecies isolated from NICUs, with respect to clinically relevant phenotypes such as-antimicrobial susceptibility, structure of the *ica* operon and biofilm formation (5, 6). Studies report increased prevalence of multidrug resistance (MDR), biofilm formation ability, and variation in the *ica* operon in *S. capitis* ssp. *ureolyticus* compared to *S. capitis* ssp. *capitis* (5), as well as differences in transcriptional response to erythromycin (6). Other published studies have also described the *S. capitis* subspecies though, none have sought to genotypically characterise them using WGS.

Most reports on *S. capitis* link it to a range of human diseases, being frequently isolated from prosthetic joint infections (7–9), prosthetic valve endocarditis (10–12) and late-onset sepsis in newborns at neonatal intensive care units (5, 13–16). The role of *S. capitis* in these infections was studied with reference to the well-described and clinically important species within the Epidermidis cluster group, i.e., *Staphylococcus epidermidis* and *Staphylococcus caprae* (17–19). *S. capitis* encodes important virulence factors required for biofilm production, persistence and immune evasion (17). *S. capitis* together with other CoNS species, such as *S. epidermidis,* are considered to have a lower virulence potential compared with the more virulent species *S. aureus,* since CoNS do not encode such an extensive suite of exotoxins associated with invasive infection (20, 21).

Species of the CoNS possess a sufficient number of virulence factors to be classed as opportunistic pathogens, contributing significantly to nosocomial infection (22). Several CoNS possess a large repertoire of virulence factor genes including those linked to adhesion and biofilm formation, affording them both commensal and pathogenic traits (20, 22, 23). CoNS are proposed to act as a reservoir for the uptake of mobile genetic elements. As commensals, they exist closely with other bacteria on skin and mucosal surfaces, and via recombination and frequent acquisition of mobile genetic elements they are able to increase their genetic diversity (23–25).

In this present study, we whole-genome sequenced 59 *S. capitis* isolates sampled from scalp skin and performed whole-genome sequencing analysis (WGSA), incorporating a further 127 publicly available sequences, with the aim to expand knowledge of *S. capitis* population structure, genotypic definition of subspecies and identify factors that are likely to contribute to virulence, competition, and colonisation.

## Materials and Methods

### Dataset and bacterial isolates

Whole-genome sequencing (WGS) was performed on 59 *S. capitis* isolates scalp skin, collected in the UK in 2017. Scalp samples were obtained using the method of Williamson and Kligman (26). The collection sample site was located and an appropriate clear and straight parting in the hair ~10 cm long was secured to maximize exposure of the scalp. A Teflon cup (18 mm internal diameter and 6 cm high) was placed onto the hair parting. A volume of 2.0 mL sampling collection medium (phosphate buffered saline plus 0.1% Triton-X-100) was applied to the collection site and the skin agitated in the liquid for one minute with a Teflon rod. The resulting 4.0 mL sample was transferred to a sterile tube. For each scalp sample taken, 100 μL of wash was plated and when possible four distinct colonies were isolated from the following staphylococcal selective medium: (1% (w/v) Tryptone (Oxoid), 0.5% (w/v) lab lemco powder (Oxoid), 0.3% (w/v) yeast extract, 1.3% (w/v) agar (Lab M), 0.1% (w/v) sodium pyruvate (JT Baker Chemicals), 0.05% (w/v) glycine (JT Baker Chemicals), 2.25% (w/v) potassium thiocyanate (JT Baker Chemicals), 0.12% (w/v) NaH2 PO4.2H20 (JT Baker Chemicals), 0.2% (w/v) lithium chloride (JT Baker Chemicals), 0.5 % (v/v) glycerol (JT Baker Chemicals), 1.0% (v/v) sodium azide (Sigma Aldrich), 3.0% (v/v) sterile egg yolk emulsion (Lab M), pH 7.2).

Additionally, 7 skin isolates, including culture collection type strains represented both *S. capitis* subspecies were sequenced and included (ATCC 49325 (*S. capitis* subsp. *ureolyticus*) and ATCC 27840 (*S. capitis* subsp. *capitis*). Also included were published genomic data available from GenBank (https://www.ncbi.nlm.nih.gov/genbank/), Sequence Read Archive (https://www.ncbi.nlm.nih.gov/sra) or European Nucleotide Archive (https://www.ebi.ac.uk/ena), and accession numbers are listed in Supplementary Table 1. All publically available *S. capitis* genomes were downloaded, subject to quality control, phylogenetically reconstructed (as described below) and Treemmer v0.3 was used to reduce redundancies within the public dataset, leaving 120 genomes to be included in further analyses (27).

### Genome sequencing and phylogenetic analysis

All isolates selectively obtained in this study, together with 7 further strains comprising type strains and skin isolates, were grown in 10 mL BHI broth (Lab M) overnight, shaking at 37 °C. Subsequently, 1 mL of each overnight culture was centrifuged for 1 min at 5000 rpm and resuspended in 180 μl lysis buffer (20 mM Tris-HCl pH8, 2 mM EDTA, 1.2% Triton X-100); the cells of each clone were extracted to obtain high-quality genomic DNA using the QIAGEN DNeasy Blood & Tissue Kit and eluted in 10 mM Tris-HCl (pH 8.5).

DNA concentration was measured using a Thermo Scientific Nanodrop, a Qubit plus visualisation after gel electrophoresis on 1% (w/v) agarose gels (at 90 mV for 40 min with a 1 kb ladder). For sequencing, the extracted DNA was required in a final volume of 60 μl with a concentration between 1-30 ng μl^-1^. Illumina DNA sequencing was performed by MicrobesNG (http://www.microbesng.uk, which is supported by the BBSRC (grant number BB/L024209/1)) using the Nextera XT library preparation kit protocol on a HiSeq platform, generating 250 bp paired end reads (Illumina, San Diego, CA, USA). The resulting datasets are available from the SRA under BioProject number PRJEB47273. Genome sequences *de novo* assembled and annotated using Unicycler v 0.4.7 (28) with default parameters and Prokka v 1.14.6 (29).

Sequence reads from 186 *S. capitis* isolates (scalp isolates, n = 59, contextual isolates, n = 127) were mapped to the reference genome *S. capitis* AYP1020 (NCBI Genome accession GCA_001028645.1) using Snippy v.4.4.3 with minimum coverage of 4 to generate core genome SNP alignment files (https://github.com/tseemann/snippy). The phylogeny of the strains was reconstructed by generating a Maximum-likelihood (ML) tree with the substitution model GTR+G+ASC and 1000 bootstrap replicates using IQTREE v 1.6.12 (30), based on the core genome alignment without the recombining regions identified by Gubbins v2.3.4 (31). Gubbins was run with default parameters on the core genome alignment of 186 strain, which generated a a chromosomal SNP alignment length of 59,972 bp. Additionally, regions containing phage identified using PHASTER (32) and MGE sequences were identified from the reference genomes, and co-ordinates were used to mask these sites, using BEDTools v 2.29.2 (33). The population structure was investigated using hierarchical Bayesian analysis of population structure with r-hierBAPS, specifying two clusters levels, 20 initial clusters and infinite extra rounds (34). Visualisations were performed using iTOL v4.2 (35).

To measure the extent of *S. capitis* genomic diversity, pairwise SNP distance was determined from the alignment of core genome SNPs identified outside regions of recombination using snp-dists v 0.7.0 (https://github.com/tseemann/snp-dists). To examine the phenotypic basis for the separation of the two subspecies within the phylogenetic reconstruction (Figure 2 & 3) the API *Staph-Ident* Stripe system was used to analyse the biochemical profiles of multiple isolates included in this study (BioMérieux, Marcy l’Etoile, France). The API Staph-Ident Strip system was used according to the manufacturer’s instructions.

### Pangenome analysis

Pangenome analysis of all 186 isolates was performed using the Panaroo v1.1.2 software package ran with default parameters, MAFFT alignment and a core gene threshold of 90% (36). Predicted coding gene sequences in all isolates were extracted, separated into core and accessory gene groups and input into eggnog-mapper v 2.1.6 to identify cluster of orthologs groups (COG) (37).

To identify genes enriched in *S. capitis* ssp. *ureolytics,* the output files from Panaroo were used as an input for Scoary v1.6.16 (38), a microbial pan-GWAS tool, to infer genes that are overrepresented in the subspecies. Scoary was ran with the –-no_pairwise flag and only genes with a Benjamini–Hochberg *p* < 0.05 and an odds ratio (OR) > 1 were considered to be overrepresented in the subspecies cluster.

### *In silico* analysis of dataset

Genetic determinants for AMR were identified using ABRicate v0.9.8 (https://github.com/tseemann/abricate) with, NCBI Bacterial Antimicrobial Resistance Reference Gene Database (39). Other potential virulence factors, such as phenol soluble modulins and exoenzymes, as well as cell wall associated proteins and two-component systems were identified by homology searches, using BLASTp of annotated reference genomes *(S. capitis* AYP1020 (Genbank assembly accession: GCA_001028645.1), *S. epidermidis* RP62a (Genbank assembly accession: GCA_000011925.1) and *S. aureus* MW2 (Genbank assembly accession: GCA_00307695.1) and pangenome data from various studies (17, 40–43).

Average nucleotide identity (ANI) indices were used to quantify genetic relatedness of the two *S. capitis* subspecies. ANI estimates the genetic relatedness between two genomes to assess species boundaries. To compare the relatedness of the *S. capitis* subspecies in this study, FastANI v1.2 was used with default settings to compare all isolates of potential ssp. *ureolyticus* to ssp. *capitis* isolates, using the recommended cut-off score of >95% that would indicate that isolates belong to the same species (44). *S. capitis* ssp. *ureolyticus* culture collection isolates were also compared to other ssp. *ureolyticus* isolates, similarly to Bannerman and Kloos (1991).

### Ethics approval

Written informed consent was obtained from all participants. The study was reviewed and approved by the Independent Ethics Committee of the Unilever Port Sunlight, United Kingdom and it was conducted in compliance with the Declaration of Helsinki and its subsequent revisions.

## Results

### Genome composition

*S. capitis* sequence reads (scalp isolates, *n* = 59, contextual isolates, *n* = 127) were assembled into draft genomes with an average of 85 contigs. The average size of the assembled genomes ranged from 2.2-2.6 Mb. Each genome had on average 2,335 (2,087– 2,565) predicted protein sequences (CDSs) with an average GC content of 32.7%.

The pan-genome of the 186 *S. capitis* isolate dataset comprised 4,471 unique clusters of orthologous groups (COGs). The pan-genome was further divided into 1,595 (35.6%) core genes (shared by all genomes) and 2,876 (64.3%) accessory genes (shared by ~95% of genomes). Gene accumulation curves reflected an open pan-genome, where the addition of each new genome increases the total gene pool (Figure 1).

**Figure 1.**
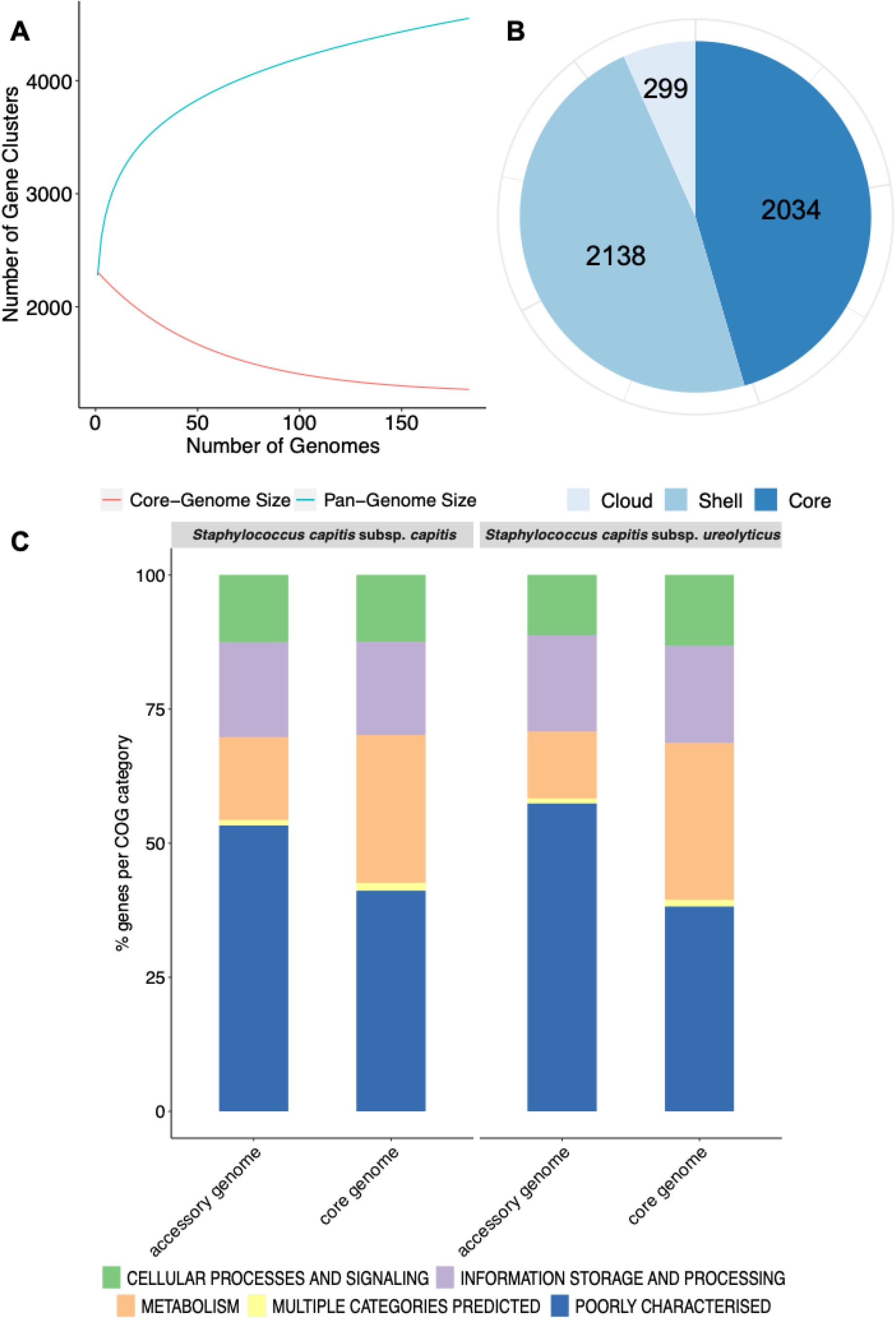
Representation of the pangenome and COG functional annotation of *S. capitis* genomes. **(A)** Pangenome curve generated by plotting total number of gene families in the pan (blue) and core (red) genome of *S. capitis.* The graph represents how the pan- and core-genomes vary as genomes are added in random order. As the number of genomes increased, the pan-genome increased. **(B)** *Staphylococcus capitis* pangenome statistics. The size of the pangenome, including core (shared by >95 % of isolates), shell (found in 15-95 % of isolates) and cloud (found in <15 % of isolates) genes. **(C)** Functional annotation of the core and accessory genomes of *S. capitis* subsp. *capitis* and *S. capitis* subsp. *ureolyticus.* Percentages of the core and accessory genomes annotated according to COG functional categories.

**Figure 2.**
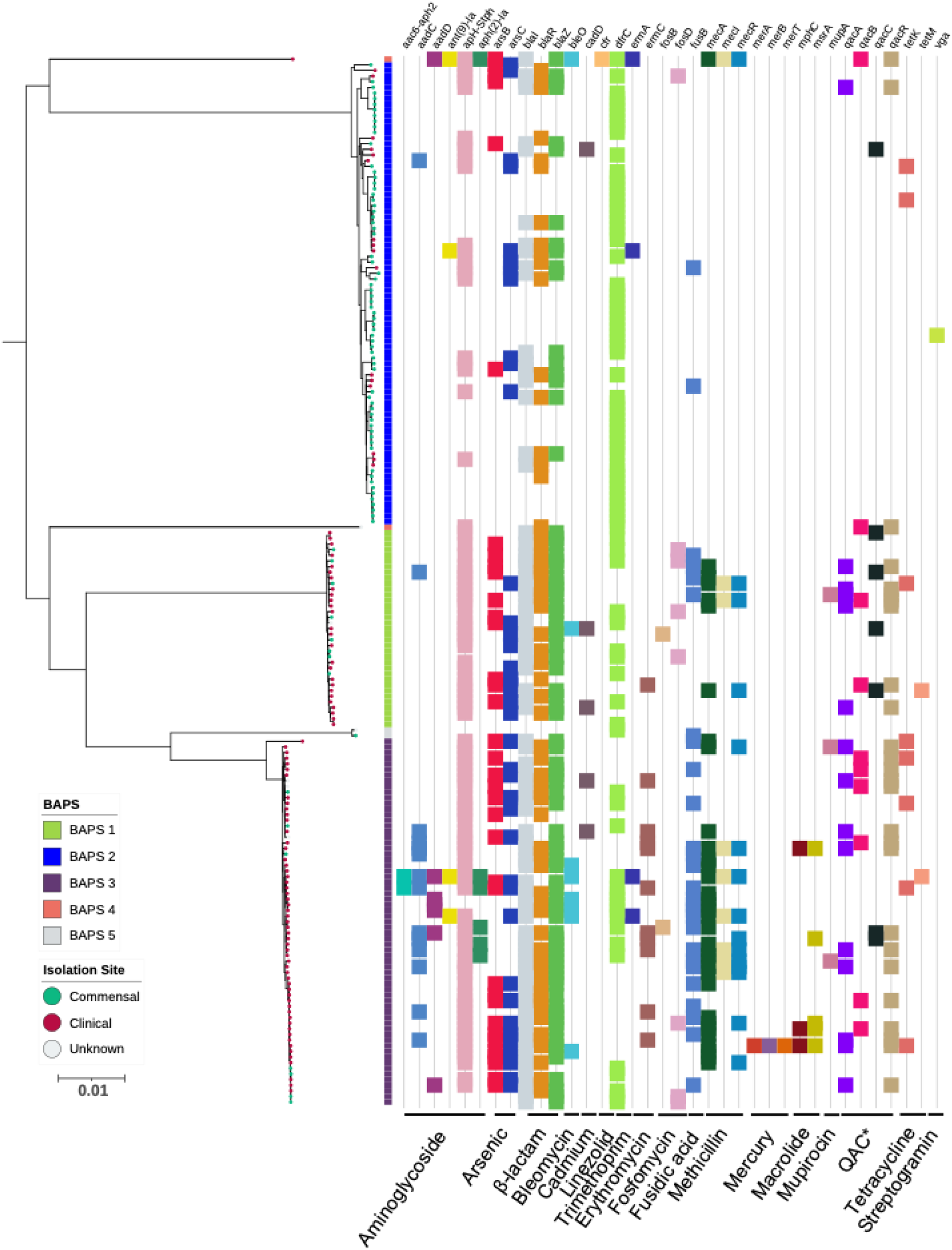
Maximum-likelihood phylogeny based on core genome alignments of 186 *S. capitis* isolates, presenting the presence and absence of antimicrobial resistance genes. ML tree is midpoint rooted and bootstrap support values were calculated from 1,000 replicates. The first colour block represents rhierBAPS clustering, dots describe the setting where isolates were retrieved; green = commensal (including scalp samples from this study), red = clinical and grey = unknown and filled grey triangles describe scalp isolates from this study. The subspecies differentiation of *S. capitis* is presented as the subclades described as BAPS groups 1, 3, 4 and 5. Presence (coloured blocks) and absence (white blocks) of antimicrobial resistance is denoted for each isolate (*Quaternary Ammonium Compounds). The scale bar represents the number of nucleotide substitutions per site. Figure was visualised using iTol v 4.2 (35).

**Figure 3.**
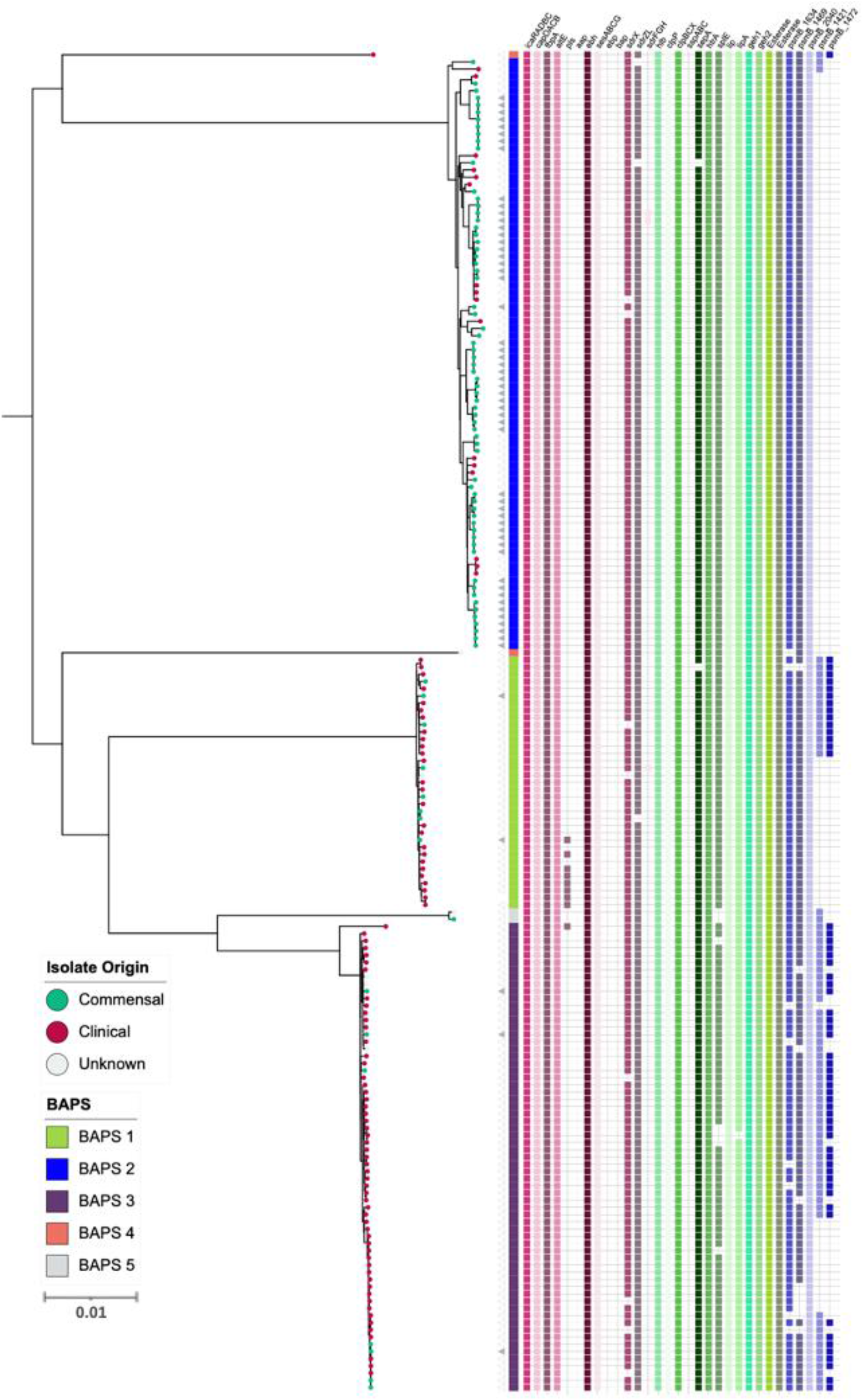
Maximum-likelihood phylogeny based on core genome alignments of 186 *S. capitis* isolates, presenting the presence and absence of genes linked to CoNS virulence potential. ML tree is midpoint rooted and bootstrap support values were calculated from 1,000 replicates. The first colour block represents rhierBAPS clustering, dots describe the setting where isolates were retrieved; green = commensal (including scalp samples from this study), red = clinical and grey = unknown and filled grey triangles describe scalp isolates from this study. The subspecies differentiation of *S. capitis* is presented as the subclades described as BAPS groups 1, 3, 4 and 5. Presence (coloured blocks) and absence (white blocks) of virulence genes is denoted for each isolate. The scale bar represents the number of nucleotide substitutions per site. Figure was visualised using iTol v 4.2 (35).

The majority of annotated genes in the accessory genomes of both *S. capitis* ssp. *capitis* (53 %) and ssp. *ureolyticus* (57 %) were poorly characterized, this was also true of the core genome (ssp. *capitis* 41 % and ssp. *ureolyticus* 38 %). This could indicate the presence of novel gene clusters in the *S. capitis* genome and the level of curation. The most abundant categories in both subspecies core genomes were those linked to essential gene function such as transcription (ssp. *capitis* 7 % and ssp. *ureolyticus* 7 %). This is in contrast to the accessory genome where gene clusters associated with replication, recombination and repair (ssp. *capitis* 9 % and ssp. *ureolyticus* 9 %) were enriched. This same pattern was separately observed in the pan-genome analysis of each subspecies (Figure 1).

### Population Structure and genetic diversity

To explore the population structure of *S. capitis,* a maximum-likelihood phylogenetic tree was constructed based on chromosomal SNP alignment length of 59,972 bp. This revealed two distinct clades separated by an average pairwise distance of 7,538 core genome SNPs, with culture collection type and contextual strains of *S. capitis* ssp. *capitis* and *S. capitis* ssp. *ureolyticus* belonging to the upper and bottom clade, respectively (Figure 2). While the *S. capitis* ssp. *capitis* clade comprised of a single dominant subclade, *S. capitis* ssp. *ureolyticus* is more diverse and comprised of 3 clades. Population structure was also inferred using BAPS to cluster genomes based on shared patterns of variation, which was congruent with the phylogeny. In this study, clinical isolates are defined as those isolated from a clinical setting e.g. hospital neonatal unit or from a disease state host. Whereas commensal isolates are defined as those isolated from healthy hosts and the community. Isolate origins were overlaid onto the phylogeny and revealed clinical isolates were predominantly found in the *S. capitis* ssp. *ureolyticus* clade, while commensal isolates were associated with the *S. capitis* ssp. *capitis* clade. However, 22 clinical isolates were observed to be interspersed across the dominant sub-clade within the *S. capitis* ssp. *capitis* clade and 77 commensal isolates were interspersed across two sub-clades within the *S. capitis* ssp. *ureolyticus* (Figure 2). This suggests that commensal and clinical isolates belonging to each subspecies are genetically similar and evolved from a common ancestor.

Extensive recombination was observed among the study isolates, with recombination most evident in BAPS clusters 2, 3, 4 and 5, which collectively contain 293 recombination blocks. BAPS cluster 1 revealed less recombination (total of 2 recombination blocks), however this is likely due to the reference genome itself being clustered in this group. Recombination was inferred across large regions of the chromosome, predominantly within the first ~750 kb and the last ~1 Mb of the genome (Figure S1). The recombination data is consistent with *S. capitis* having arisen following extensive recombination events (13).

### Insights into *S. capitis* pathogenicity

The CoNS that colonise human skin are generally considered to be non-pathogenic species specialised for healthy human skin and mucosal surfaces, but they are now emerging as important opportunistic pathogens (20, 22, 24, 45–47). Antimicrobial resistance properties are important factors of nosocomial infection. Therefore, we screened for presence of genetic determinants for AMR and identified genes predicted to encode for resistance against tetracyline, β-lactam, bleomycin, fosfomycin, methicillin resistance, fusidic acid streptogramin A, macrolide, linezolid, trimethoprim and aminoglycoside. Amongst the *S. capitis* genomes analysed, 48 % (5 % *S. capitis* ssp. *capitis* and 44 % *S. capitis* ssp. *ureolyticus*) were classified as MDR, carrying genetic determinants conferring resistance against three or more classes of drugs (Figure 2). MDR was found in 78 % of isolates from the *ureolyticus* clade, compared to 11 % of isolates from the *capitis* clade. When the dataset was stratified by isolate origin, we found clinical isolates carried more AMR genes than compared to commensal. A total of 635 AMR linked genes were found across clinical isolates (*n* = 99) compared to 169 found in commensal isolates (*n* = 81).

In addition to AMR genes, we also investigated the role of phenol-souble modulins (PSMs) contributing to the virulence potential of *S. capitis.* PSMs are a novel toxin family that have antimicrobial activity (48, 49) and have been attributed to the competitive success of CoNS due to their ability to inhibit the growth of other commensal bacteria such as *Cutibacterium acnes* (O’Neill et al., 2020). A total of 5 gene clusters encoding β-class PSMs were identified (Figure 3), with gene clusters 1634, 1469 and 2040 found in >98 % of isolates, sharing >90 % similarity when locally aligned to PSMs described and isolated from *S. capitis,* by O’Neill *et al.,* (O’Neill et al., 2020). Similar to AMR gene presence and absence, PSM-associated gene clusters were found more abundantly in the *S. capitis* ssp. *ureolyticus* clade than compared with the *S. capitis* ssp. *capitis* clade. Specifically, gene clusters 1421 and 1474 were found in 66 % and 55 % of isolates from the *ureolyticus* clade, in contrast to 3 % and 0 % isolates from the *capitis* clade. (Figure 4).

**Figure 4.**
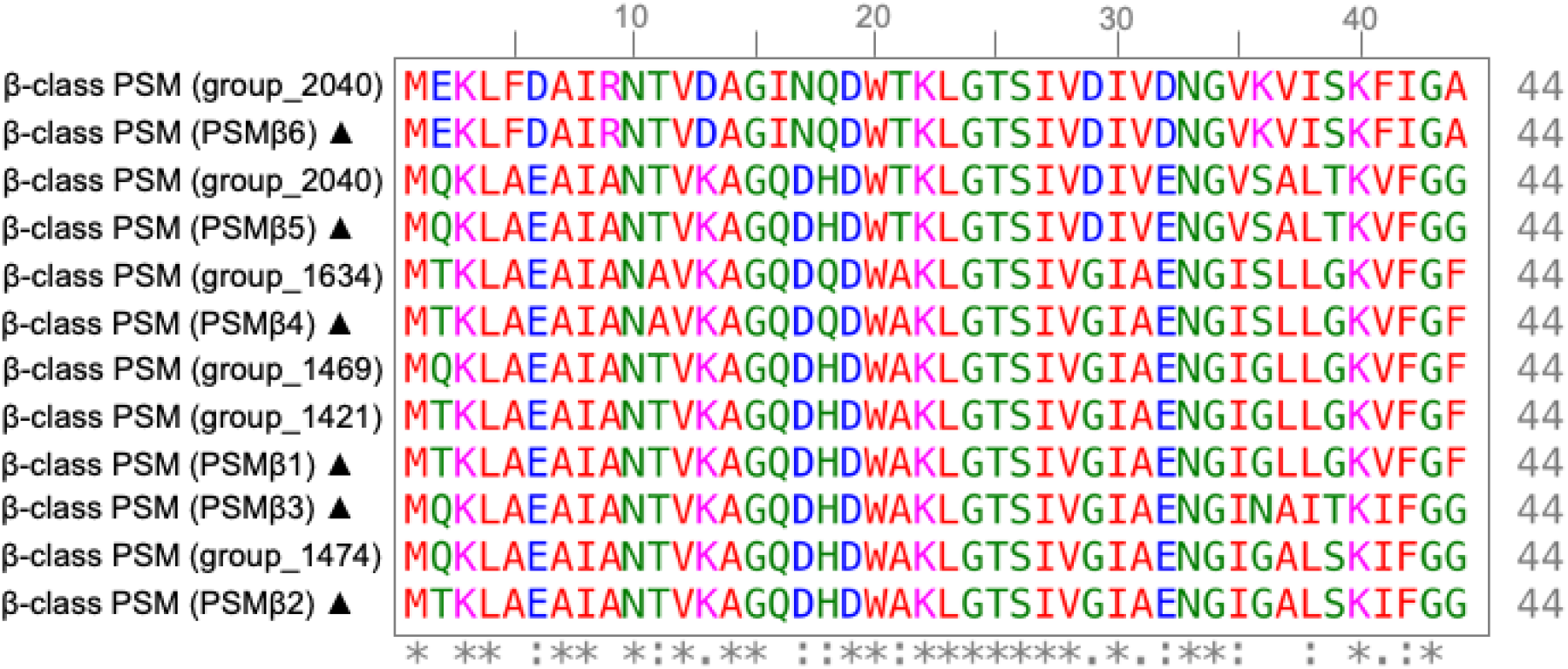
Multiple sequence alignment of β-class Phenol Soluble Modulins (PSMs) of *S. capitis* isolates. MSA of β-class PSMs protein sequences found in *S. capitis* genomes from this study and those described by O’Neill *et al.,* (2020) (sequences marked with ▲) created with ClustalW (65). Residues are coloured based on amino acid property (Red: small and hydrophobic, blue: acidic, magenta: basic, green: hydroxyl, sulfhydryl, and amine and grey: unusual), positions that contain fully conserved residues are marked with and asterisk and positions marked with a colon indicate conservation between groups of amino acids with similar properties.

To further investigate variation of the β-class PSMs identified, we performed multiple sequence alignment of corresponding amino acid sequence from the 6 gene clusters, which revealed conservation of residues attributed to the maintenance of the amphipathic nature of the peptides, essential to PSM antimicrobial activity (50). Specifically, lysine at 3^rd^ and/or tryptophan at 20^th^ position are putatively associated with providing antibacterial activity of β-class family peptides (50), both of which are conserved in the peptide sequences of this study (Figure 4).

Additionally, we specifically screened for the presence of orthologous CDS that likely contribute to *S. capitis* pathogenicity. Including, staphylococcal cell wall associated (CWA) proteins curated with potential virulence roles, including biofilm-associated proteins IcaRADBC, capsule biosynthesis proteins CapDACB, surface adhesins AtlE, Pls, Aap, FnbpA, SesA, SesB, SesC, SesG, Ebp and Bap, and MSCRAMMs SdrX, SdrZL, SdrH, SdrF and SdrG. Of these CWA proteins, 11 were encoded in *S. capitis* AYP1020. Across the 186 *S. capitis* genomes analysed, *sesA, sesB, sesC, sesG, icaRADBC, fbnpA, capDACB* and *atlE* were found in all isolates. MSCRAMM genes *sdrX* and *sdrZL* were found in >95 % of genomes investigated. Contrastingly, the surface adhesin gene *pls* was found in <10 % of *S. capitis* genomes and was absent from *ssp. capitis* (Figure 3). Notably, genes that were absent from the *S. capitis* AYP1020 genome (determined in this study as *S. capitis* ssp. *ureolyticus)* but present in other CoNS species such as, *S. epidermidis* RP62a (Cameron et al., 2015), included *sdrF, ebp, bap, sdrH, sdrG* and *aap.* These genes were absent from >95 % of the genomes included in this study (17) (Figure 3). The secreted protein genes *hlb, clpP, clpBCX, sepA, htrA, lip, geh1, geh2* and *lipA* were present in >95 % of all the isolates. The presence of a suite of exoproteins could contribute to host colonisation, persistence, infection and immune evasion, important to both pathogenesis and colonisation (22). Notably, *sspA, sspB* and *sspC* were absent in *S. capitis* genomes, the serine protease SspA promotes invasion in *S. aureus* (51) (Figure 3).

The 16 two component systems (TCS) described in *S. aureus* are conserved in other closely related CoNS species, indicative of an adaptive and highly versatile pathogen (52, 53). *S. aureus* TCS are extensively characterised and therefore the 186 *S. capitis* genomes included in this study were screened for homologous protein sequences. Of the 16 TCS, 14 were found in *S. capitis.* This is intermediate with *S. epidermidis* (16 TCS) and *S. saprophyticus* (11 TCS) with number determined by Rapun-Araiz, et al. (2020) to be indeterminate of genome size in staphylococci (Table 1).

**Table 1.** Two-component systems in *S. aureus* and *S. capitis.* Presence and absence of the 16 TCS of *S. aureus* (MW2) described in *S. capitis* reference genome AYP1020 and isolates included in this study.

| Two-component system | *S. aureus* reference | *S. capitis* reference | Presence in *S. capitis*<br>ssp. *capitis* (%) | Presence in *S. capitis*<br>ssp. *ureolyticus* (%) | Function |
| --- | --- | --- | --- | --- | --- |
| *walRK* | MW0018 | AYP1020_RS09955 | 100 | 100 | Cell wall maintenance, cell viability |
|  | MW0019 | AYP1020_RS09960 | 100 | 100 |  |
| *hptSR* | MW0198 |  | 0 | 0 | Intracellular survival, uptake of hexose phosphate |
|  | MW0199 |  | 0 | 0 |  |
| *lytSR* | MW0236 |  | 0 | 0 | Autolysis, eDNA release, biofilm |
|  | MW0237 |  | 0 | 0 |  |
| *graRS* | MW0621 | AYP1020_RS00130 | 100 | 100 | AMPs resistance, growth at low pH |
|  | MW0622 | AYP1020_RS00135 | 100 | 100 |  |
| *saeSR* | MW0667 | AYP1020_RS00365 | 100 | 100 | Virulence factors regulation |
|  | MW0668 | AYP1020_RS00360 | 100 | 100 |  |
| *tcs7SR* | MW1208 | AYP1020_RS03075 | 100 | 100 | Uncharacterised function |
|  | MW1209 | AYP1020_RS03080 | 100 | 100 |  |
| *arlRS* | MW1304 | AYP1020_RS03545 | 100 | 100 | Pathogenesis mechanisms |
|  | MW1305 | AYP1020_RS03540 | 100 | 100 |  |
| *srrBA* | MW1445 | AYP1020_RS03875 | 100 | 100 | Anaerobic respiration, metabolism, growth at low temperatures |
|  | MW1446 | AYP1020_RS03880 | 100 | 100 |  |
| *phoRP* | MW1636 | AYP1020_RS04760 | 100 | 100 | Phosphate uptake and homeostasis |
|  | MW1637 | AYP1020_RS04765 | 100 | 100 |  |
| *airRS* | MW1789 | AYP1020_RS05370 | 100 | 100 | Oxidative stress response |
|  | MW1790 | AYP1020_RS05375 | 100 | 100 |  |
| *varRS* | MW1824 | AYP1020_RS05700 | 100 | 100 | Cell wall-affecting antibiotic resistance, cell wall biosynthesis |
|  | MW1825 | AYP1020_RS05705 | 100 | 100 |  |
| *agrCA* | MW1962 | AYP1020_RS06010 | 100 | 100 | Quorum sensing control of adhesion and virulence factors |
|  | MW1963 | AYP1020_RS06005 | 100 | 100 |  |
| *kpdDE* | MW2002 | AYP1020_RS09655 | 100 | 100 | Potassium homeostasis regulation |
|  | MW2003 | AYP1020_RS09660 | 100 | 100 |  |
| *hssRS* | MW2282 | AYP1020_RS07580 | 100 | 100 | Heme metabolism regulation |
|  | MW2283 | AYP1020_RS07585 | 100 | 100 |  |
| *nreCB* | MW2313 | AYP1020_RS07750 | 100 | 100 | Response to low oxygen, nitrate reduction |
|  | MW2314 | AYP1020_RS07755 | 100 | 100 |  |
| *braSR* | MW2544 | AYP1020_RS08920 | 100 | 100 | Antimicrobial peptide resistance |
|  | MW2545 | AYP1020_RS08925 | 100 | 100 |  |

### *S. capitis* subspecies definition

To biochemically assess differences between the two subspecies *S. capitis* ssp. *capitis* and *S. capitis* ssp. *ureolyticus* based upon the original descriptions of Bannerman and Kloos (2) the API *Staph-Ident* Strip system was used (BioMérieux, Marcy l’Etoile, France). A total of 22 isolates sampled in this study that were spread across the phylogenetic tree (11 assigned to the *S. capitis* ssp. *capitis* clade and 11 assigned to the *S. capitis* ssp. *ureolyticus* clade), as well as type and culture collection stains were tested for classifying phenotypic traits. Among the 11 isolates belonging to the *S. capitis* ssp. *ureolyticus* clade, only 4 (33 %) tested urease-positive/maltose-positive and 10 (91%) *S. capitis* ssp. *capitis* isolate tested urease-negative/maltose-negative, indicating an unreliable phenotype of urease activity and maltose fermentation as a subspecies definition (data not shown).

Since the original phenotypic trait descriptors for *S. capitis* did not sufficiently discriminate between the subspecies, we then sought to quantify and analyse the genetic differences between *S. capitis* ssp. *ureolyticus* and *capitis.* Analysis of the average nucleotide identity (ANI) between isolates determined here as *S. capitis ssp. ureolyticus* and *S. capitis ssp. capitis* revealed that genomes from the two subspecies shared little genetic differences with 96% ANI. Pan-genomic comparative analysis also revealed limited gene content variation of 1% between the two subspecies. To help identify genes that could be used to discriminate between the subspecies and serve as diagnostic markers for rapid identification by PCR in future studies, we looked for genes which are significantly overrespresented in each subspecies and identified total of 38 gene clusters found across all *S. capitis ssp. capitis* genomes and 13 across all *S. capitis ssp. ureolyticus* genomes (Table 2, Supplementary Table 2 & 3). Upon closer inspection, majority of the subspecies unique genes identified were in fact divergent gene orthologues. An example of this is the *icaC* gene, which encodes for an intercellular adhesion protein. While one version of this gene cluster was only found in *S. capitis* ssp. *ureolyticus* isolates (98 %), another version sharing blastp identity of 22 % was only present in *S. capitis* ssp. *capitis* isolates (0 %).

**Table 2.** Gene clusters found significantly enriched in either *S. capitis* ssp. *capitis* or ssp. *ureolyticus* (*p* < 0.001). Gene clusters, presence and absence, and functional descriptions were obtained from Panaroo and Scoary pangenome analysis of assembled genomes. X these genes exist in different conserved versions in isolates. *S. capitis* reference gene numbers are from *S. capitis* AYP1020 (Genbank Assembly Accession: GCA_001028645.1) (17). The complete list can be found at Supplementary Table 2.

| Gene name | *S. capitis* reference | Annotation | *S.capitis* ssp. *capitis* (%) | *S.capitis* ssp. *ureolyticus* (%) |
| --- | --- | --- | --- | --- |
| group_2869 |  | Hypothetical protein | 100 | 0 |
| group_459 |  | Putative phage head morphogenesis protein | 100 | 0 |
| crtNX |  | Dehydrosqualene desaturase | 100 | 4.76 |
| hdcA |  | Histidine decarboxylase proenzyme | 100 | 6.67 |
| hdsM |  | Type I restriction-modification system methyltransferase subunit | 100 | 21.90 |
| group_1023 |  | Putative transcriptional regulator | 100 | 29.52 |
| group_2665 | AYP1020_RS09385 | Putative protein YjdF | 100 | 34.29 |
| sasCX |  | Cell-wall-anchored protein SasC | 100 | 35.24 |
| arcAX |  | Arginine deiminase | 98.72 | 6.67 |
| arcCX |  | Carbamate kinase ArcC1 | 98.72 | 7.62 |
| arcDX |  | Arginine/ornithine APC family amino acid-polyamine-organocation transporter antiporter | 98.72 | 7.62 |
| argFX |  | Ornithine carbamoyltransferase | 98.72 | 7.62 |
| hdsSX |  | Type I restriction modification DNA specificity protein;type I restriction modification system site specificity determination subunit HsdS_1 | 98.72 | 7.62 |
| trkG |  | Trk family potassium (K+) transporter ABC protein | 98.72 | 11.43 |
| arsB |  | Arsenical pump membrane protein | 98.72 | 16.19 |
| cdr |  | Coenzyme A disulfide reductase | 98.72 | 19.05 |
| arsR |  | Arsenical resistance operon repressor | 98.72 | 19.05 |
| group_2407 |  | Arsenate reductase (glutaredoxin) | 98.72 | 20.95 |
| arsA |  | Arsenical pump-driving ATPase | 98.72 | 55.24 |
| arsD |  | Arsenical resistance operon trans-acting repressor ArsD | 92.31 | 37.14 |
| pyrBX |  | Aspartate carbamoyltransferase catalytic subunit | 98.72 | 0.95 |
| group_1353 | AYP1020_RS02690 | Hypothetical protein | 0 | 100 |
| group_798 | AYP1020_RS11575 | Hypothetical protein | 0 | 100 |
| group_1413 | AYP1020_RS11570 | Hypothetical protein | 0 | 99.05 |
| icaCX | AYP1020_RS08445 | Putative poly-beta-16-N-acetyl-D-glucosamine export protein | 0 | 98.10 |
| fmrO | AYP1020_RS12300 | rRNA methyltransferase FmrO;hypothetical protein | 35.90 | 96.19 |
| dapF | AYP1020_RS09635 | Diaminopimelate epimerase | 35.90 | 96.19 |
| qorB | AYP1020_RS06480 | Cobalt (Co2+) ABC superfamily ATP binding cassette transporter membrane protein | 0 | 78.10 |
| group_1472 |  | Antibacterial protein (phenol soluble modulin) | 0 | 55.24 |
| mqo |  | Putative malate:quinone oxidoreductase 2 | 0 | 64.76 |
| pfpI |  | Intracellular protease | 0 | 60 |
| opp1B |  | Oligopeptide ABC superfamily ATP binding cassette transporter membrane protein | 35.90 | 84.76 |
| opp1D |  | Oligopeptide ABC superfamily ATP binding cassette transporter ABC protein | 35.90 | 84.76 |
| group_687 |  | Hypothetical protein | 35.90 | 84.76 |
| group_585 |  | MFS family major facilitator transporter | 35.90 | 84.76 |
| opp1A |  | Oligopeptide ABC superfamily ATP binding cassette transporter binding protein | 35.90 | 84.76 |
| opp1C |  | Oligopeptide ABC superfamily ATP binding cassette transporter membrane protein | 35.90 | 84.76 |
| opp1F |  | ABC superfamily ATP binding cassette transporter ABC protein | 35.90 | 84.76 |
| blaRX | | β-lactamase regulatory protein | 6.41 | 62.86 |

To further investigate specific genetic signatures associated with each *S. capitis* subspecies, we applied Scoary to identify genes that are overrepresented in *S. capitis* ssp. *capitis,* as well as *S. capitis* ssp. *ureolyticus.* A total of 1,086 predicted gene clusters were found to differ significantly (*p* < 0.05) between the subspecies, although there were no significant differences found between assigned functional COG categories (Table 1, Supplementary Table 2, Figure 1). Gene clusters with a known function identified as being enriched in *S. capitis* ssp. *capitis* isolates include those for the arginine catabolic mobile element (ACME), encoding arginine deiminase activity found in various species of staphylococci. Most *S. capitis* isolates in this study contain a Type V ACME gene cluster, however different conserved versions are found in each subspecies. Additional gene clusters enriched in ssp. *capitis* included those encoding: SasC cell wall anchored protein; and CrtN dehydrosqualene desaturase involved in staphyloxanthin biosynthesis (Table 2, Supplementary Table 2 & 3). For *S. capitis* ssp. *ureolyticus,* genes predicted to encode for virulence factor were found to be enriched. Specifically, genes with antimicrobial-associated functions including β-lactam resistance protein BlaR and β-class phenol soluble modulins (Table 2, Supplementary Table 2 & 3). This is concurrent with the current literature that *S. capitis* ssp. *ureolyticus* as the more virulent subspecies (5, 8).

## Discussion

*S. capitis* is an opportunistic pathogen that is associated with increasing reports of bloodstream infections and neonatal infections in intensive care units. Currently, *S. capitis* is mostly studied with reference to the well-described and clinically important *S. epidermidis*. In the absence of an expansive study of *S. capitis* genomes from the scalp, the current work aimed to explore WGS of the species to expand knowledge of its population structure, compare and contrast genomic differences between commensal and clinical isolates to gain understanding of the genetic factors that contributes to *S. capitis* pathogenicity.

Pangenome analysis indicated that the *S. capitis* has an open pangenome, which could arise from a capacity to acquire exogenous DNA whilst living in extensive bacterial communities. The large accessory genome size suggests that *S. capitis* contains a large repertoire of genes that confer advantages under particular environmental conditions to support its colonisation and/or cause infections in clinical settings. Similarly, other members of the Epidermidis cluster group, such as *S. caprae* and *S. epidermidis* also have a large, open pangenome state compared to other CoNS-*Staphyloccus lugeunensis* (54, 55). It can therefore be hypothesised that *S. capitis,* like other staphylococci, undergo horizontal gene transfer (HGT) which could also have led to the acquisition of virulence genes within *S. capitis* genomes (54). The identification of more virulent strains of *S. capitis* ssp. *ureolyticus,* in greater frequencies in clinical settings combined with less virulent strains isolated from other sources, such as the scalp could indicate a potential contextual basis for the HGT events, although further investigation is needed to understand the association of *S. capitis* subspecies and scalp skin state. Many studies using the higher resolution afforded by WGS have enabled important differences to be uncovered in *S. capitis* genomes, such as their multidrug resistance profiles across different geographic regions (5, 13, 17).

Phylogenetic analysis revealed clustering of two distinct clades that likely represent the two subspecies, herein termed the *S. capitis* spp. *capitis* and the *S. capitis* spp. *ureolyticus* clade. Most of the clinical isolates from this study belonged to the *S. capitis* spp. *ureolyticus* clade and commensal isolates belonging to the *S. capitis* spp. *capitis* clade, suggesting that *S. capitis* spp. *ureolyticus* is more associated with clinical infections. This agrees with the current literature that suggests *S. capitis* spp. *ureolyticus* is the more virulent subspecies and linked to the presence of genes linked to biofilm formation and methicillin resistance (5, 8). However, the observation of clinical isolates interspaced across the dominant sub-clade of *S. capitis* spp. *capitis* clade indicates that clinical and commensal isolates share similar genetic background, and while *S. capitis* spp. *capitis* is more associated with commensal it can cause clinical infection.

Although the scalp-associated isolates sampled in this study were not from clinical infections, potential virulence-linked genes were found throughout their genomes, highlighting *S. capitis* versatility and potential for adaptation that might cause significant disease in settings like the NICU. Further exploration of sequence differences will be required to unravel defining features of *S. capitis* subspecies to support the hypothesis that *S. capitis* ssp. *ureolyticus* or genomes belonging to the *S. capitis* ssp. *ureolyticus* clade, are generally more virulent.

Most published studies have focused on *S. capitis* strains isolated from NICU and other clinical settings, describing the emergence of drug resistance in response to the use of antimicrobial and antiseptic therapy to treat infections caused by CoNS isolates (13–15, 56, 57). Antimicrobial resistance to vancomycin and fusidic acid has been reported among *S. capitis,* like *S. aureus,* suggesting the occurrence of inter-species genetic exchange (13, 56).

In addition to antimicrobial resistance, biofilm formation was proposed to be an important virulence trait of *S. capitis* in both a clinical and commensal setting (5, 20). In keeping with this proposal, the current work confirms the presence of biofilm-related genes in the *S. capitis* genomes studied, including *icaRADBC* operon, *ebh* and *atlE,* and extends it by identifying an IcaC encoding gene cluster discriminating the subspecies with its presence in ssp. *ureolyticus.* The role of IcaC (Table 2) in *S. capitis* has been described by Cui, et al as modifying synthesised glucan by acetylation (2015). A more extensive investigation, into IcaC, including *S. capitis* isolated from different sources could contribute to further understanding trait differences that determine ssp. specialisation. The suite of biofilm genes facilitate staphylococcal primary attachment by binding to extracellular matrix molecules, and intercellular aggregation (20). The presence of these genes may confer a selective advantage in both a clinical setting and on the scalp and forehead. Further phenotypic studies of biofilm formation, metabolism, and multidrug resistance in *S. capitis* isolates, including those in this study will extend our knowledge of speciation and specialisation of staphylococci and *S. capitis,* and could help with precise studies of factors that pertain to emergent clinical disease.

Currently, subdivision of *S. capitis* to ssp. *ureolyticus* and ssp. *capitis* is based upon original descriptions of *S. capitis* ssp. *ureolyticus* urease activity, ability to produce acid from maltose, fatty acid profile, larger colony size and DNA sequence differentiation (2). The multiple discriminating traits were not explored in full here but biochemical analysis using API *Staph-Ident* Strip system, the most common method to discriminate *S. capitis* subspecies, was tested on 14 isolates sampled from this study and 3 isolates from the culture collection type strains. Discrepancy between biochemical and whole genome phylogenetic assignment of the subspecies were observed, as only 33 % of tested isolates that belonged to the *S. capitis* ssp. *ureolyticus* clade tested positive for urease activity. Highlighting that classification of *S. capitis* subspecies by urease activity is unreliable and requires confirmation using other discriminating traits.

Further analysis to characterise the genetic relatedness between the two subspecies using ANI revealed genomes of each subspecies were similar, sharing 96 % nucleotide identity. Instead, we investigated discriminating gene clusters and observed that *S. capitis* spp. *ureolyticus* genomes were enriched with antimicrobial resistance gene functions, such as β-lactam resistance genes and β-class phenol soluble modulins. Whereas *S. capitis* spp. *capitis* genomes were enriched with gene clusters linked to skin survival based on the presence of the arginine catabolism mobile element (ACME) that encodes enzymes to counteract low pH (58, 59). ACME is a genomic island first described in *S. aureus* USA300 and in *S. epidermidis* ATCC 12228 (60, 61). It was shown to enhance staphylococcal colonisation of the skin and mucous membranes, showing similar characteristics to the staphylococcal cassette chromosome *mec* (SCC*mec*) element (62, 63). While not investigated functionally here in *S. capitis* it is likely to have a similar function. We hypothesise that ACME activity, that discriminates ssp. *capitis,* could represent a key factor of subspeciation through niche specialisation. Analysis of virulence gene profiles determined that as a species, *S. capitis* has a similar repertoire of virulence genes to several other CoNS species, with respect to AMR, PSMs and secreted proteases (54, 64). Investigation of the subspecies classifications highlighted here demonstrate that further analysis is required for robust markers of subspecies classification within their core genomes though several genes exclusive to every isolate of each subspecies were identified and could serve as subspecies biomarkers.

In conclusion, this study identified distinct clustering of the two subspecies of *S. capitis* and determined gene clusters for traits that might rapidly progress our understanding of *S. capitis* relevant to disease. Specifically, we propose that the original subspecies definition ssp. *ureolyticus* needs reconsidered based on species subclades that define it based on the importance of MDR and virulence. It is likely that the widespread use of antimicrobials, the openness of the *S. capitis* pangenome and acquisition of MGEs with beneficial mutations has promoted the emergence of virulence traits in *S. capitis* isolates. Continued research into classification of *S. capitis* as subspecies versus virulence linked phylogroups is important for better surveillance of the spread and evolution of *S. capitis.*

## Data Summary

Short read sequences supporting the findings of this study have been deposited in the European Nucleotide Archive (https://www.ebi.ac.uk/ena/) under the project accession number PRJEB47273. Accession numbers for isolates used in this study are listed in Supplementary Table 1. Publicly available sequences were downloaded from GenBank (https://www.ncbi.nlm.nih.gov/genbank/), Sequence Read Archive (https://www.ncbi.nlm.nih.gov/sra) or European Nucleotide Archive (https://www.ebi.ac.uk/ena), and accession numbers are listed in Table Supplementary Table 1.

## Acknowledgments

DNA sequencing was performed by the Centre for Genomic Research, Liverpool, UK. We are grateful to Sally Grimshaw for technical assistance and Barry Murphy for technical discussions.

## Author Notes

C.E.C performed all the data analysis and interpretation for the results under the scientific guidance of R.J.B and M.J.H. C.E.C and M.J.H drafted the manuscript. All authors contributed to editing of the manuscript.

All supporting data, code and protocols have been provided within the article or through supplementary data files. Three supplementary tables and one supplementary figure are available.

## Supplementary Information

**Figure S1. Analysis of the *S. capitis* genome alignment with Gubbins.** The maximum likelihood phylogenetic reconstruction of *S. capitis* is shown on the left, with coloured bands highlighting rhierBAPS clustering and isolation site of genomes. Filled grey triangles describe scalp isolates from this study. Homologous recombination events for each *S. capitis* genome ordered based on their position in the AYP1020 reference genome (shown along the top) are shown on the right. Recombination blocks detected in >1 isolate are shown in red, while blocks affecting a single isolate are indicated in blue. Figure visualised using Phandango (66).

**Table S1. *Staphylococcus capitis* isolates used in this study.** Isolates were either isolated in this study, type strains or sequence data obtained from the NCBI database.

**Table S2. Genes significantly differentiating *S. capitis* subspecies.** Gene identification output from Scoary analysis.

**Table S3. Representative protein sequences of hypothetical gene clusters that significantly differentiate *S. capitis* subspecies.** Gene identification output from Scoary analysis and protein sequence information from annotated genome assemblies.

## References

1. Kloos WE, Schleifer KH. Isolation and characterization of staphylococci from human skin II. descriptions of four new species: Staphylococcus warneri, Staphylococcus capitis, Staphylococcus hominis, and Staphylococcus simulans1. International Journal of Systematic and Evolutionary Microbiology. 1975;25(1):62–79.

2. Bannerman TL, Kloos WE. Staphylococcus capitis subsp. ureolyticus subsp. nov. from human skin. Int J Syst Bacteriol. 1991;41(1):144–7.

3. Maggs AF, Pennington TH. Temporal study of staphylococcal species on the skin of human subjects in isolation and clonal analysis of Staphylococcus capitis by sodium dodecyl sulfate-polyacrylamide gel electrophoresis. J Clin Microbiol. 1989;27(12):2627–32.

4. Grimshaw SG, Smith AM, Arnold DS, Xu E, Hoptroff M, Murphy B. The diversity and abundance of fungi and bacteria on the healthy and dandruff affected human scalp. PLoS One. 2019;14(12):e0225796.

5. Cui B, Smooker PM, Rouch DA, Daley AJ, Deighton MA. Differences between two clinical Staphylococcus capitis subspecies as revealed by biofilm, antibiotic resistance, and pulsed-field gel electrophoresis profiling. Journal of clinical microbiology. 2013;51(1):9–14.

6. Cui B, Smooker PM, Rouch DA, Deighton MA. Effects of erythromycin on the phenotypic and genotypic biofilm expression in two clinical Staphylococcus capitis subspecies and a functional analysis of Ica proteins in S. capitis. Journal of medical microbiology. 2015;64(6):591–604.

7. Tevell S, Hellmark B, Nilsdotter-Augustinsson Å, Söderquist B. Staphylococcus capitis isolated from prosthetic joint infections. Eur J Clin Microbiol Infect Dis. 2017;36(1):115–22.

8. Tevell S, Baig S, Hellmark B, Martins Simoes P, Wirth T, Butin M, et al. Presence of the neonatal Staphylococcus capitis outbreak clone (NRCS-A) in prosthetic joint infections. Scientific Reports. 2020;10(1):22389.

9. Flurin L, Greenwood-Quaintance KE, Patel R. Microbiology of polymicrobial prosthetic joint infection. Diagnostic Microbiology and Infectious Disease. 2019;94(3):255–9.

10. Nalmas S, Bishburg E, Meurillio J, Khoobiar S, Cohen M. Staphylococcus capitis prosthetic valve endocarditis: Report of two rare cases and review of literature. Heart & Lung. 2008;37(5):380–4.

11. Cone LA, Sontz EM, Wilson JW, Mitruka SN. Staphylococcus capitis endocarditis due to a transvenous endocardial pacemaker infection: Case report and review of Staphylococcus capitis endocarditis. International Journal of Infectious Diseases. 2005;9(6):335–9.

12. Al Hennawi HET, Mahdi EM, Memish ZA. Native valve Staphylococcus capitis infective endocarditis: a mini review. Infection. 2020;48(1):3–5.

13. Carter GP, Ussher JE, Da Silva AG, Baines SL, Heffernan H, Riley TV, et al. Genomic Analysis of Multiresistant Staphylococcus capitis Associated with Neonatal Sepsis. Antimicrobial agents and chemotherapy. 2018;62(11):e00898–18.

14. Wirth T, Bergot M, Rasigade J-P, Pichon B, Barbier M, Martins-Simoes P, et al. Niche specialization and spread of Staphylococcus capitis involved in neonatal sepsis. Nature Microbiology. 2020;5(5):735–45.

15. Rasigade J-P, Raulin O, Picaud J-C, Tellini C, Bes M, Grando J, et al. Methicillin-Resistant Staphylococcus capitis with Reduced Vancomycin Susceptibility Causes Late-Onset Sepsis in Intensive Care Neonates. PLoS One. 2012;7(2):e31548.

16. Stenmark B, Hellmark B, Söderquist B. Genomic analysis of Staphylococcus capitis isolated from blood cultures in neonates at a neonatal intensive care unit in Sweden. European Journal of Clinical Microbiology & Infectious Diseases. 2019;38(11):2069–75.

17. Cameron D, Jiang J-H, Hassan K, Elbourne L, Tuck K, Paulsen I, et al. Insights on virulence from the complete genome of Staphylococcus capitis. Frontiers in Microbiology. 2015;6(980).

18. Lamers RP, Muthukrishnan G, Castoe TA, Tafur S, Cole AM, Parkinson CL. Phylogenetic relationships among Staphylococcus species and refinement of cluster groups based on multilocus data. BMC Evol Biol. 2012;12:171.

19. Watanabe S, Aiba Y, Tan X-E, Li F-Y, Boonsiri T, Thitiananpakorn K, et al. Complete genome sequencing of three human clinical isolates of Staphylococcus caprae reveals virulence factors similar to those of S. epidermidis and S. capitis. BMC Genomics. 2018;19(1):810.

20. Otto M. Staphylococcus epidermidis — the ‘accidental’ pathogen. Nature Reviews Microbiology. 2009;7(8):555–67.

21. Massey RC, Horsburgh MJ, Lina G, Höök M, Recker M. The evolution and maintenance of virulence in Staphylococcus aureus: a role for host-to-host transmission? Nature Reviews Microbiology. 2006;4(12):953–8.

22. Heilmann C, Ziebuhr W, Becker K. Are coagulase-negative staphylococci virulent? Clinical Microbiology and Infection. 2019;25(9):1071–80.

23. Becker K, Heilmann C, Peters G. Coagulase-Negative Staphylococci. Clinical Microbiology Reviews. 2014;27(4):870.

24. Otto M. Coagulase-negative staphylococci as reservoirs of genes facilitating MRSA infection: Staphylococcal commensal species such as Staphylococcus epidermidis are being recognized as important sources of genes promoting MRSA colonization and virulence. Bioessays. 2013;35(1):4–11.

25. Argemi X, Hansmann Y, Prola K, Prévost G. Coagulase-Negative Staphylococci Pathogenomics. International journal of molecular sciences. 2019;20(5):1215.

26. Williamson P, Kligman AM. A new method for the quantitative investigation of cutaneous bacteria. The Journal of investigative dermatology. 1965;45(6):498–503.

27. Menardo F, Loiseau C, Brites D, Coscolla M, Gygli SM, Rutaihwa LK, et al. Treemmer: a tool to reduce large phylogenetic datasets with minimal loss of diversity. BMC Bioinformatics. 2018;19(1):164.

28. Wick RR, Judd LM, Gorrie CL, Holt KE. Unicycler: Resolving bacterial genome assemblies from short and long sequencing reads. PLoS computational biology. 2017;13(6):e1005595–e.

29. Seemann T. Prokka: rapid prokaryotic genome annotation. Bioinformatics. 2014;30(14):2068–9.

30. Chernomor O, von Haeseler A, Minh BQ. Terrace Aware Data Structure for Phylogenomic Inference from Supermatrices. Systematic Biology. 2016;65(6):997–1008.

31. Croucher NJ, Page AJ, Connor TR, Delaney AJ, Keane JA, Bentley SD, et al. Rapid phylogenetic analysis of large samples of recombinant bacterial whole genome sequences using Gubbins. Nucleic Acids Research. 2014;43(3):e15–e.

32. Arndt D, Grant JR, Marcu A, Sajed T, Pon A, Liang Y, et al. PHASTER: a better, faster version of the PHAST phage search tool. Nucleic Acids Res. 2016;44(W1):W16–21.

33. Quinlan AR, Hall IM. BEDTools: a flexible suite of utilities for comparing genomic features. Bioinformatics. 2010;26(6):841–2.

34. Cheng L, Connor TR, Sirén J, Aanensen DM, Corander J. Hierarchical and Spatially Explicit Clustering of DNA Sequences with BAPS Software. Molecular Biology and Evolution. 2013;30(5):1224–8.

35. Letunic I, Bork P. Interactive tree of life (iTOL) v3: an online tool for the display and annotation of phylogenetic and other trees. Nucleic Acids Res. 2016;44(W1):W242–5.

36. Tonkin-Hill G, MacAlasdair N, Ruis C, Weimann A, Horesh G, Lees JA, et al. Producing polished prokaryotic pangenomes with the Panaroo pipeline. Genome Biology. 2020;21(1):180.

37. Huerta-Cepas J, Forslund K, Coelho LP, Szklarczyk D, Jensen LJ, von Mering C, et al. Fast Genome-Wide Functional Annotation through Orthology Assignment by eggNOG-Mapper. Mol Biol Evol. 2017;34(8):2115–22.

38. Brynildsrud O, Bohlin J, Scheffer L, Eldholm V. Rapid scoring of genes in microbial pan-genome-wide association studies with Scoary. Genome biology. 2016;17(1):1–9.

39. Feldgarden M, Brover V, Haft DH, Prasad AB, Slotta DJ, Tolstoy I, et al. Validating the AMRFinder Tool and Resistance Gene Database by Using Antimicrobial Resistance Genotype-Phenotype Correlations in a Collection of Isolates. Antimicrobial agents and chemotherapy. 2019;63(11):e00483–19.

40. Gill Steven R, Fouts Derrick E, Archer Gordon L, Mongodin Emmanuel F, DeBoy Robert T, Ravel J, et al. Insights on Evolution of Virulence and Resistance from the Complete Genome Analysis of an Early Methicillin-Resistant Staphylococcus aureus Strain and a Biofilm-Producing Methicillin-Resistant Staphylococcus epidermidis Strain. Journal of Bacteriology. 2005;187(7):2426–38.

41. Baba T, Bae T, Schneewind O, Takeuchi F, Hiramatsu K. Genome sequence of Staphylococcus aureus strain newman and comparative analysis of staphylococcal genomes: Polymorphism and evolution of two major pathogenicity islands. Journal of Bacteriology. 2008;190(1):300–10.

42. Baba T, Takeuchi F, Kuroda M, Yuzawa H, Aoki K-i, Oguchi A, et al. Genome and virulence determinants of high virulence community-acquired MRSA. The Lancet. 2002;359(9320):1819–27.

43. Altschul SF, Gish W, Miller W, Myers EW, Lipman DJ. Basic local alignment search tool. J Mol Biol. 1990;215(3):403–10.

44. Jain C, Rodriguez-R LM, Phillippy AM, Konstantinidis KT, Aluru S. High throughput ANI analysis of 90K prokaryotic genomes reveals clear species boundaries. Nature Communications. 2018;9(1):5114.

45. Post V, Harris LG, Morgenstern M, Mageiros L, Hitchings MD, Méric G, et al. Comparative Genomics Study of Staphylococcus epidermidis Isolates from Orthopedic-Device-Related Infections Correlated with Patient Outcome. Journal of clinical microbiology. 2017;55(10):3089–103.

46. Méric G, Mageiros L, Pensar J, Laabei M, Yahara K, Pascoe B, et al. Disease-associated genotypes of the commensal skin bacterium Staphylococcus epidermidis. Nature Communications. 2018;9(1):5034.

47. Espadinha D, Sobral RG, Mendes CI, Méric G, Sheppard SK, Carriço JA, et al. Distinct Phenotypic and Genomic Signatures Underlie Contrasting Pathogenic Potential of Staphylococcus epidermidis Clonal Lineages. Frontiers in Microbiology. 2019;10(1971).

48. Cheung GYC, Joo H-S, Chatterjee SS, Otto M. Phenol-soluble modulins –critical determinants of staphylococcal virulence. FEMS Microbiology Reviews. 2014;38(4):698–719.

49. Peschel A, Otto M. Phenol-soluble modulins and staphylococcal infection. Nature Reviews Microbiology. 2013;11(10):667–73.

50. Kumar R, Jangir PK, Das J, Taneja B, Sharma R. Genome Analysis of Staphylococcus capitis TE8 Reveals Repertoire of Antimicrobial Peptides and Adaptation Strategies for Growth on Human Skin. Scientific reports. 2017;7(1):10447–.

51. Dubin G, Chmiel D, Mak P, Rakwalska M, Rzychon M, Dubin A. Molecular cloning and biochemical characterisation of proteases from Staphylococcus epidermidis. Biological chemistry. 2001;382(11):1575–82.

52. Haag AF, Bagnoli F. The Role of Two-Component Signal Transduction Systems in Staphylococcus aureus Virulence Regulation. In: Bagnoli F, Rappuoli R, Grandi G, editors. Staphylococcus aureus: Microbiology, Pathology, Immunology, Therapy and Prophylaxis. Cham: Springer International Publishing; 2017. p. 145–98.

53. Rapun-Araiz B, Haag AF, Solano C, Lasa I. The impact of two-component sensorial network in staphylococcal speciation. Current Opinion in Microbiology. 2020;55:40–7.

54. Sun Z, Zhou D, Zhang X, Li Q, Lin H, Lu W, et al. Determining the Genetic Characteristics of Resistance and Virulence of the “Epidermidis Cluster Group” Through Pan-Genome Analysis. Front Cell Infect Microbiol. 2020;10:274–.

55. Argemi X, Matelska D, Ginalski K, Riegel P, Hansmann Y, Bloom J, et al. Comparative genomic analysis of Staphylococcus lugdunensis shows a closed pan-genome and multiple barriers to horizontal gene transfer. BMC genomics. 2018;19(1):621–.

56. Chong J, Caya C, Lévesque S, Quach C. Heteroresistant Vancomycin Intermediate Coagulase Negative Staphylococcus in the NICU: A Systematic Review. PLoS One. 2016;11(10):e0164136.

57. Mello D, Daley AJ, Rahman MS, Qu Y, Garland S, Pearce C, et al. Vancomycin Heteroresistance in Bloodstream Isolates of <em>Staphylococcus capitis</em&gt. Journal of Clinical Microbiology. 2008;46(9):3124.

58. Lindgren JK, Thomas VC, Olson ME, Chaudhari SS, Nuxoll AS, Schaeffer CR, et al. Arginine Deiminase in Staphylococcus epidermidis Functions To Augment Biofilm Maturation through pH Homeostasis. Journal of Bacteriology. 2014;196(12):2277–89.

59. Planet PJ, LaRussa SJ, Dana A, Smith H, Xu A, Ryan C, et al. Emergence of the Epidemic Methicillin-Resistant Staphylococcus aureus Strain USA300 Coincides with Horizontal Transfer of the Arginine Catabolic Mobile Element and <i>speG</i>-mediated Adaptations for Survival on Skin. mBio. 2013;4(6):e00889–13.

60. Planet PJ, Diaz L, Kolokotronis S-O, Narechania A, Reyes J, Xing G, et al. Parallel Epidemics of Community-Associated Methicillin-Resistant Staphylococcus aureus USA300 Infection in North and South America. The Journal of Infectious Diseases. 2015;212(12):1874–82.

61. Diep BA, Gill SR, Chang RF, Phan TH, Chen JH, Davidson MG, et al. Complete genome sequence of USA300, an epidemic clone of community-acquired meticillin-resistant Staphylococcus aureus. The Lancet. 2006;367(9512):731–9.

62. Diep BA, Stone GG, Basuino L, Graber CJ, Miller A, des Etages S-A, et al. The Arginine Catabolic Mobile Element and Staphylococcal Chromosomal Cassette mec Linkage: Convergence of Virulence and Resistance in the USA300 Clone of Methicillin-Resistant Staphylococcus aureus. The Journal of Infectious Diseases. 2008;197(11):1523–30.

63. Almebairik N, Zamudio R, Ironside C, Joshi C, Ralph JD, Roberts AP, et al. Genomic Stability of Composite SCCmec ACME and COMER-Like Genetic Elements in Staphylococcus epidermidis Correlates With Rate of Excision. Frontiers in Microbiology. 2020;11(166).

64. Moawad AA, Hotzel H, Awad O, Roesler U, Hafez HM, Tomaso H, et al. Evolution of Antibiotic Resistance of Coagulase-Negative Staphylococci Isolated from Healthy Turkeys in Egypt: First Report of Linezolid Resistance. Microorganisms. 2019;7(10):476.

65. Thompson JD, Higgins DG, Gibson TJ. CLUSTAL W: improving the sensitivity of progressive multiple sequence alignment through sequence weighting, position-specific gap penalties and weight matrix choice. Nucleic Acids Res. 1994;22(22):4673–80.

66. Hadfield J, Croucher NJ, Goater RJ, Abudahab K, Aanensen DM, Harris SR. Phandango: an interactive viewer for bacterial population genomics. Bioinformatics. 2017;34(2):292–3.

