## Supplementary figures and images for "Comparative genomics of *Staphylococcus capitis* reveals determinants of speciation"

### Figure S1

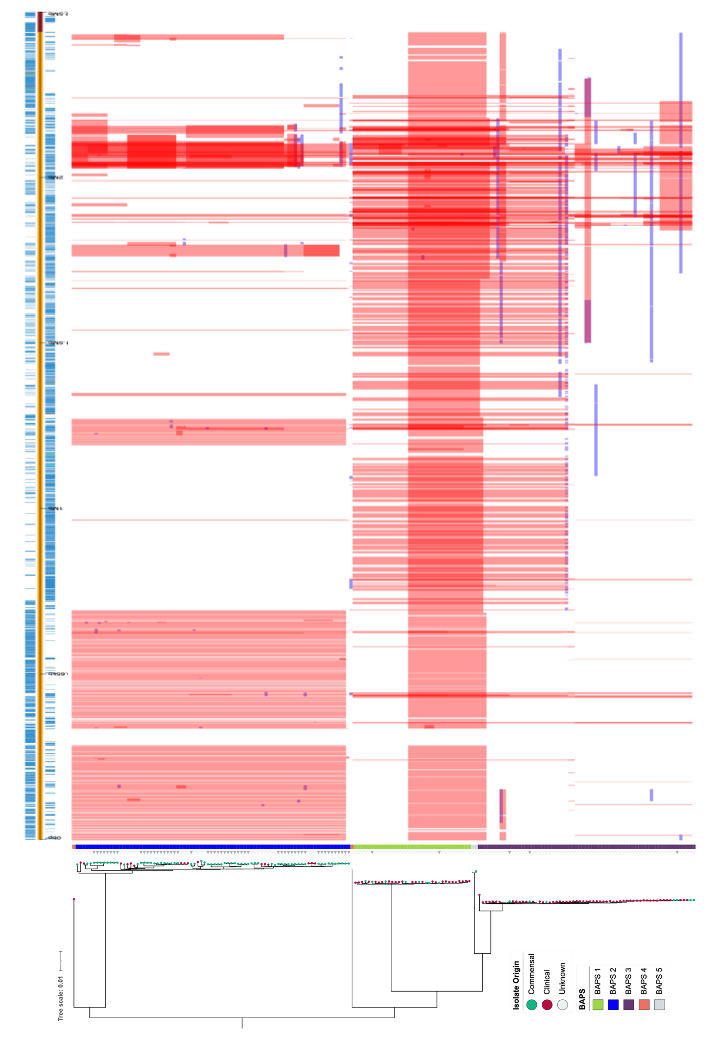
